# CORnet: Modeling the Neural Mechanisms of Core Object Recognition

**DOI:** 10.1101/408385

**Authors:** Jonas Kubilius, Martin Schrimpf, Aran Nayebi, Daniel Bear, Daniel L. K. Yamins, James J. DiCarlo

## Abstract

Deep artificial neural networks with spatially repeated processing (a.k.a., deep convolutional ANNs) have been established as the best class of candidate models of visual processing in primate ventral visual processing stream. Over the past five years, these ANNs have evolved from a simple feedforward eight-layer architecture in AlexNet to extremely deep and branching NAS-Net architectures, demonstrating increasingly better object categorization performance and increasingly better explanatory power of both neural and behavioral responses. However, from the neuroscientist’s point of view, the relationship between such very deep architectures and the ventral visual pathway is incomplete in at least two ways. On the one hand, current state-of-the-art ANNs appear to be too complex (e.g., now over 100 levels) compared with the relatively shallow cortical hierarchy (4-8 levels), which makes it difficult to map their elements to those in the ventral visual stream and to understand what they are doing. On the other hand, current state-of-the-art ANNs appear to be not complex enough in that they lack recurrent connections and the resulting neural response dynamics that are commonplace in the ventral visual stream. Here we describe our ongoing efforts to resolve both of these issues by developing a “CORnet” family of deep neural network architectures. Rather than just seeking high object recognition performance (as the state-of-the-art ANNs above), we instead try to reduce the model family to its most important elements and then gradually build new ANNs with recurrent and skip connections while monitoring both performance and the match between each new CORnet model and a large body of primate brain and behavioral data. We report here that our current best ANN model derived from this approach (CORnet-S) is among the top models on Brain-Score, a composite benchmark for comparing models to the brain, but is simpler than other deep ANNs in terms of the number of convolutions performed along the longest path of information processing in the model. All CORnet models are available at github.com/dicarlolab/CORnet, and we plan to update this manuscript and the available models in this family as they are produced.

## Introduction

The apparently effortless human ability to rapidly recognize visually presented objects in various contexts has long fascinated neuroscientists, cognitive scientists, and computer vision researchers alike. To investigate the underpinnings of this process, DiCarlo et al. (2012) operationalized the behavioral domain *core object recognition (COR)* in which each subject must discriminate from all other possible objects a dominant object with the viewing duration of a natural fixation (∼200 ms) in the central visual field (10 deg) under high view and background variation. Over the past five years, deep artificial neural networks with repeated spatially-local processing (a.k.a. deep convolutional ANNs) that are optimized to solve COR-like tasks have emerged as the leading class of models in that they very accurately predict – for *any* image in central 10 deg – both the detailed pattern of COR behavioral responses and the neural patterns of responses at successive stages of the ventral visual stream (see Yamins and DiCarlo (2016) for an overview) that causally support COR behavior (Holmes and Gross, 1984; Horel et al., 1987; Afraz et al., 2015; Moeller et al., 2017; Rajalingham and DiCarlo, 2018).

As first shown by (Yamins et al., 2013, 2014), models in the deep convolutional ANN class whose parameters are optimized to achieve high performance tend to develop internal “neural” representations that are closest to those observed in the highest levels of the ventral visual stream (V4 and IT). While this result implies that continued discovery and optimization of even higher performing ANN models will lead to even better models of the ventral visual stream and COR behavior, that implication cannot hold forever without additional constraints from neuroscience. For example, the highest performing deep convolutional ANNs have abandoned their original tether to the approximate number of cortical processing stages in the ventral visual stream (4-8) and are now well over 100 stages in depth. While that architectural evolution has – at least in the case of large amounts of supervised trained data – led to higher performing models as judged by the current metrics of computer vision (esp. ImageNet categorization performance; Russakovsky et al., 2015), the connection to neurobiology has grown far more murky in that it is unclear which, if any, model layer(s) are putative models of specific ventral stream cortical areas (e.g., V1, V2, V4, IT). Moreover, these newer ANN models are invariably feedforward, in contrast to abundant lateral and feedback connections and their resulting response dynamics known to exist in the ventral visual stream.

We do not criticize the machine learning and computer vision communities for pursuing these less biologically-tied newer models (e.g., deeper, more supervised training data, etc.) because higher system performance is indeed an important technological goal, and gains clearly are being made by this approach, at least for the near term. However, as brain and cognitive scientists, our job is to harness the aspects of those newer models that are mostly likely to inform an understanding of the brain and cognition and, ideally, to use that deeper understanding to make further long-term gains in machine learning and computer vision.

To do that we are continuously building ANNs that: better approximate the brain architecture, adopt the latest ideas from state-of-the-art ANNs, and maintain or improve their functional match to brain and behavioral data associated with core object recognition. We refer to these ANNs as the CORnet family. In this paper, we report on our current progress and release to the community three members of the CORnet family that we hope will be useful to others. Our best model so far, CORnet-S, holds one of the current best scores a set of quantitative benchmarks for judging a model’s match to neural and behavioral responses (Brain-Score.org; Schrimpf et al., 2018) yet it is much more compact than competing networks from computer vision.

## Model criteria

### General criteria (based on Kubilius, 2018)

1. **Predictive:** Our major requirement is to have models that are the most explanatory of neural and behavioral benchmarks. We use *Brain-Score* (Schrimpf et al., 2018) to quantify this aspect of the goodness of our models.
2. **Compact:** Among model that score well on Brain-Score, we prefer the simpler ones as they are potentially easier to understand and more efficient to experiment with. We introduce the notion of *Feedforward Simplicity* as one measure of simplicity.
3. **Computable:** Ideally, a model should act just like any other participant in an experiment, receiving the same instructions and producing outputs without any free parameters left for a researcher to fine-tune. We thus prefer models that fully specify all computation details: how to go from inputs to outputs, specific weights and hyper-parameters, and so on. In other words, we expect any proposed model to be a concrete commitment that could be potentially falsified by experimental data.

### Specific criteria for Core Object Recognition models

1. **Internally mappable:** Since we are modeling the brain, we are not only interested in having correct model outputs (behaviors) but also internals that match brain’s anatomical and functional constraints. We prefer neural network models because neurons are the units of online information transmission and models without neurons cannot be obviously mapped to neural spiking data (Yamins and DiCarlo, 2016).
2. **Few layers:** The human and non-human primate ventral visual pathway consists of only a handful of areas that process visual inputs: retina, LGN, V1, V2, V4, and a set of areas in the inferior temporal cortex (IT). While the exact number of areas is not yet established, we ask that models have few areas (though each area may perform multiple operations).
3. **Single canonical circuitry in all model areas:** We have no strong reason to believe that circuitry should be different across areas in the ventral visual pathway. We therefore prefer models where the operations and connectivity in each model area are the same.
4. **Recurrent:** While core object recognition was originally believed to be the result of largely feedforward mechanism because of its fast time scale (DiCarlo et al., 2012), it has long been suspected that recurrent connections must be relevant for some aspects of object perception (Lamme and Roelfsema, 2000; Bar et al., 2006; Wagemans et al., 2012), and recent studies have shown a potential role of recurrent processes even at the fast time scale of core object recognition (Kar et al., 2018a; Tang et al., 2018; Rajaei et al., 2018). Moreover, even if recurrent processes are not critical to core object recognition, responses in the visual system still have a temporal profile, so a good model at least should be able to produce responses *over time*.

## Brief history of modeling vision

Modeling human visual processing traces back at least to Hubel and Wiesel where response properties of simple cells in visual area V1 were formalized as feature detection of edges and properties of complex cells were conceptualized as a set of operations that were spatially repeated over the visual field (Hubel and Wiesel, 1962, i.e., translationally invariant). These computational principles inspired the first models of object recognition, most notably, the Neocognitron (Fukushima, 1980) and the HMAX model family (Riesenhuber and Poggio, 1999; Serre et al., 2007), where feature detectors and pooling operators were used in turns to build deep hierarchical models of object recognition. However, such models lacked robust feature representations as it was not clear at the time how to either build in or otherwise train these networks to learn their spatially-repeated operations from input statistics – particularly for areas beyond visual area V1 (Olshausen and Field, 1996; Lowe, 1999; Torralba and Oliva, 2003). These issues were first addressed by the AlexNet ANN (Krizhevsky et al., 2012) in that it demonstrated at least one way to train a deep neural network for a large-scale invariant object recognition task (Russakovsky et al., 2015). Concurrently, deep networks optimized for such invariant object recognition tasks were demonstrated to produce internal “neural” representations that were by far the best models of the responses of neurons in non-human primate visual areas V4 and IT (Yamins et al., 2013; Cadieu et al., 2014; Yamins et al., 2014). Later work in humans confirmed these gains in explanatory power at the courser experimental level of fMRI and MEG (Khaligh-Razavi and Kriegeskorte, 2014; Güçlü and van Gerven, 2015; Cichy et al., 2016), with detailed measures of behavioral response patterns in both humans and non-human primates (e.g., Rajalingham et al., 2015; Kubilius et al., 2016; Rajalingham et al., 2018), and with non-human primate neural spiking measures from the cortical area V1 (Cadena et al., 2017).

Today, the best known models for core object recognition, as measured by *Brain-Score*, are very deep feedforward networks, such as ResNet-101 (He et al., 2016) and DenseNet-169 (Huang et al., 2017). In this work, we aimed to simplify these model classes as much as possible while maintaining their brain predictivity and meeting other requirements listed in the **Introduction** that we believe good models of the ventral stream should feature.

## CORnet family of architectures

CORnet family of architectures (Fig. 1) has four computational areas, conceptualized as analogous to the visual areas V1, V2, V4, and IT, and a linear category decoder that maps from the population of neurons in the model’s last visual area to its behavioral choices. Each visual area implements a particular neural circuitry with neurons performing simple canonical computations: convolution, addition, non-linearity, response normalization or pooling over a receptive field. In each of the three CORnet models presented here (Z, R, S), the circuitry is identical in each of its visual areas, but we vary the total number of neurons in each area. Below we describe three particular circuitries that we identified as useful for predicting brain responses (Fig. 1, bottom, and Table 1).

**Table 1.**
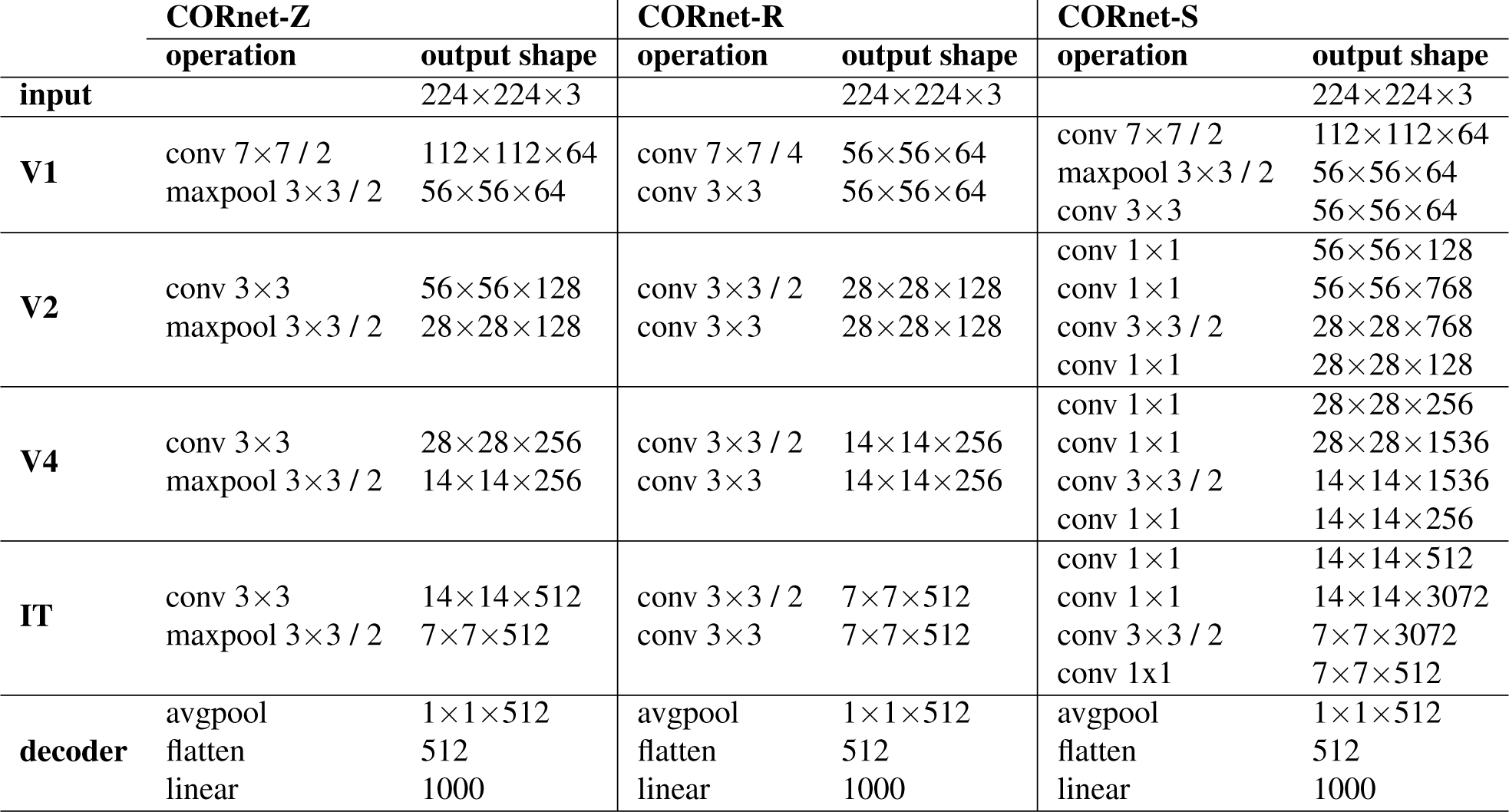
Architectural parameters of each model.

**Fig. 1.**
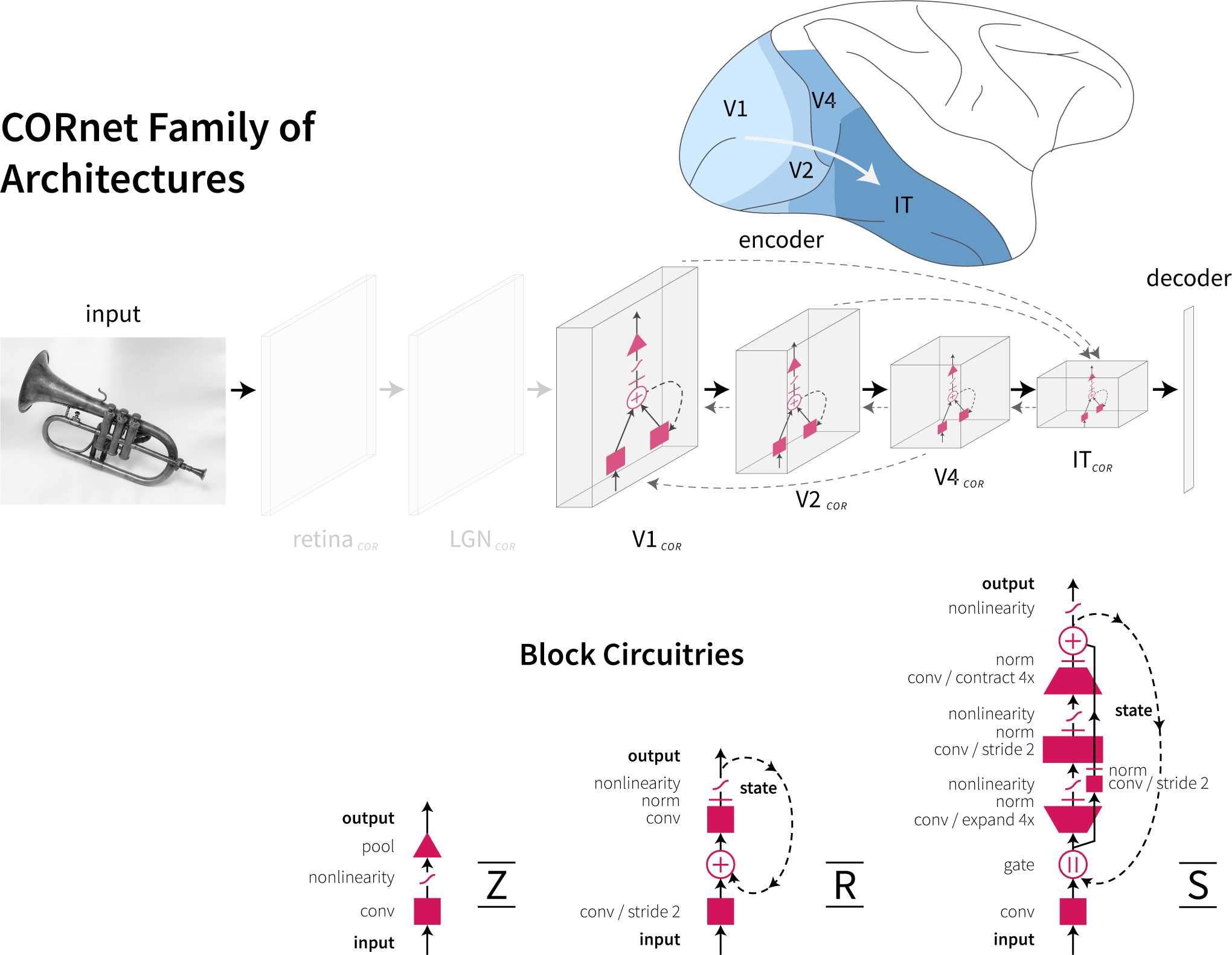
CORnet family of architectures. All models in this family have four areas that are pre-mapped to cortical areas V1, V2, V4, and IT. Retinal and LGN processing is currently not modeled *(greyed-out areas)*. All areas can be either feedforward or recurrent (within area) but currently we do not consider models with skip or feedback connections between areas *(grayed-out dashed arrows)*. In this paper, we consider three flavors of CORnet, each serving distinct functions: CORnet-Z is a lightweight AlexNet alternative, CORnet-R is a simple recurrent model with biologically-plausible unrolling in time, and CORnet-S is our highest-performing model on Brain-Score at the moment. Diagrams in the bottom panel depict circuitry within an area of each model.

The decoder part of a model implements a simple linear classifier – a set of weighted linear sums with one sum for each object category. When training on ImageNet, responses of last model area (and last time step in the case of recurrence) are further passed through a softmax nonlinearity to perform a 1000-way classification. To reduce the amount of neural responses projecting to this classifier, we first average responses over the entire receptive field per feature map

There are no across-area bypass or across-area feedback connections in the current definition of CORnet family. Retinal and LGN processing are omitted (see **Discussion**). Also, due to high computational demands, the first area in CORnet-S is simplified to only include two convolutions (see below).

### CORnet-Z

(a.k.a. “Zero”) is our simplest model, derived by observing that (i) AlexNet is already nearly as good in predicting neural responses as deeper models (Schrimpf et al., 2018) and (ii) multiple fully-connected layers do not appear necessary to achieve good ImageNet performance, as most architectures proposed after VGG (Simonyan and Zisserman, 2014) contain only a singe 1000-way linear classification layer. Thus, CORnet-Z’s area circuits consist of only a single convolution, followed by a ReLU nonlinearity and max pooling.

### CORnet-R

(a.k.a. “Recurrent”) introduces recurrent dynamics that would propagate through the network in a biologically-valid manner (see Fig. 2 for a comparison between biological and non-biologicall network unrolling). In CORnet-R, recurrence is introduced only within an area (no feedback connections between areas), so the particular way of unrolling has little effect (apart from consuming much more memory), but we chose to use biologically-valid unrolling nonetheless to make this model useful for investigating neural dynamics.

**Fig. 2.**
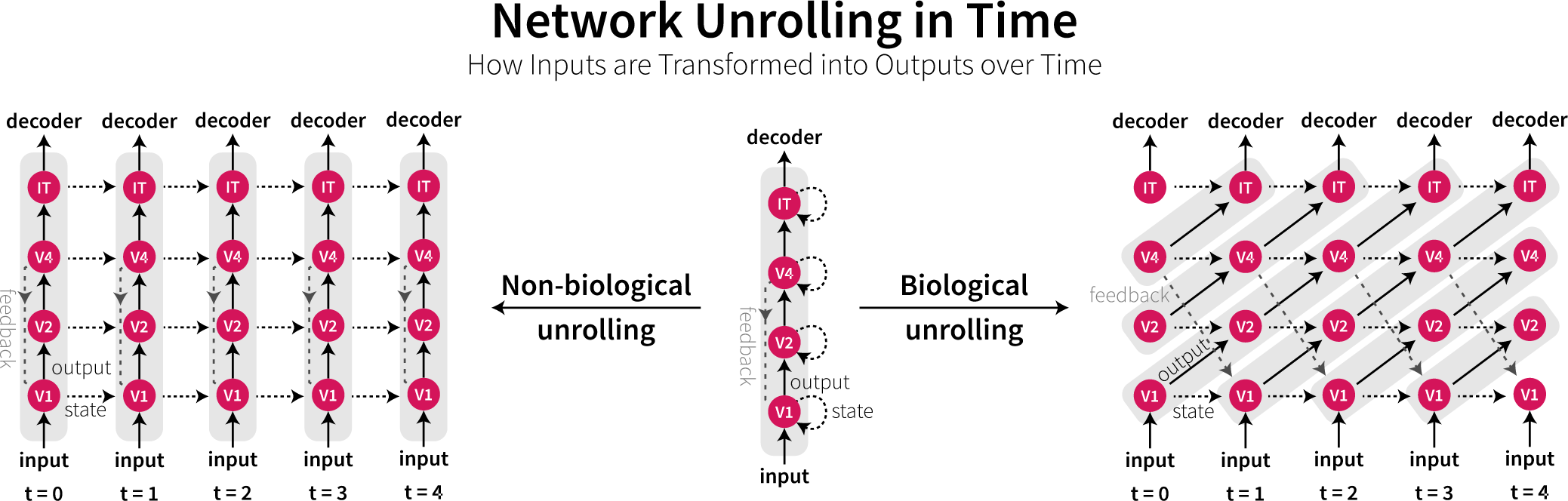
Network unrolling in time. A recurrent network needs to pass information through its computational units multiple times. This poses a problem: What is the order of computations? In standard machine learning implementations of recurrent models (*left*), information is passed at each time step from inputs to outputs (i.e., to decoder in our case). In contrast, in biological systems (*right*), inputs are transformed to outputs in a stepwise fashion. At *t* = 0, only the first area (V1 in this case) processes an input and sends the output of this computation to V2. At *t* = 1, V2 processes V1’s output, while V1 is already processing a new input with an updated internal state. It takes four time steps for the original input to finally reach IT, whereas in a non-biological unrolling of a network the input reaches IT at the same time as it is fed to V1. The diagram on the right illustrates how we apply recurrence in CORnet-R. The horizontal dashed lines are the implementation of the within area recurrence shown as dashed lines in Fig. 1. The difference between the two kinds of unrolling would become most dramatic when a feedback connection is introduced (*dim gray dashed arrows from V4 to V1*). In standard recurrent networks, information from V4 would be sent down to V1 but since V1 has already processed its inputs, this information would not be utilized. In biological systems, at *t* = 1 feedback from V4 would be combined with inputs to V1, so it would readily affect further processing downstream of V1.

The input is first downscaled twofold while increasing the number of channels twofold by passing it through a convolution, followed by normalization and a nonlinearity. The state (initially zero) is added to the result and passed through another convolution, normalization, and nonlinearity, and the result is saved as a new state of the area. We used group normalization (Wu and He, 2018) and ReLU nonlinearity.

### CORnet-S

(a.k.a. “Skip”) aims to rival the best models on Brain-Score by transforming very deep feedforward architectures into a shallow recurrent model. Specifically, CORnet-S draws inspiration from ResNets that are some of the best models on our behavioral benchmark (Rajalingham et al., 2018; Schrimpf et al., 2018). Liao and Poggio (2016) proposed that ResNet structure could be thought of as an unrolled recurrent network and recent studies further demonstrated that weight sharing was indeed possible without a significant loss in CIFAR and ImageNet performance (Jastrzęb-ski et al., 2017; Leroux et al., 2018).

CORnet-S stacks two more convolutions (each followed by a normalization and nonlinearity) on top of CORnet-R’s circuit. Moreover, following ResNet’s bottleneck block structure, the second convolution expands the number of channels fourfold while the last one decreases them back. We also included a skip connection, such that the result of adding the state to the input is combined with the output of the last convolution just prior to applying a nonlinearity. Given that CORnet-S only has within-area recurrent connections, in order to minimize memory footprint we trained this model without making use of any network unrolling in time (i.e. no unrolling as depicted in Fig. 2, but weights are still shared over repeated computations). In particular, the first time information is processed, the third convolution and the skip convolution use a stride of two to downscale inputs. The output is then used instead of the original input for further recurrent processing (referred to as “gate” in Fig. 1). We used batch normalization (Ioffe and Szegedy, 2015) and ReLU nonlinearity in this model, and batch normalization was not shared over time as suggested by (Jastrzębski et al., 2017).

### Implementation details

- **Framework:** PyTorch 0.4.1
- **Data:**
  **- Dataset:** ImageNet 2012 (Russakovsky et al., 2015)
  **- Preprocessing:**
    **\* Train:** Random crop to 224 x 224 px image, random left/right flip, mean subtraction and division by standard deviation
    **\* Validation:** Central crop to 224 x 224 px image, mean subtraction and division by standard deviation.
  **- Batch:** 256 images, trained on a single (CORnet-S and CORnet-R) NVIDIA Titan X GPU / GeForce 1080Ti or divided over 2 GPUs (CORnet-S)
  **- Training duration:** 25 epochs (CORnet-Z and CORnet-R); 43 epochs (CORnet-S)
- **Learning rate:** We use similar learning rate scheduling to ResNet with more variable learning rate updates (primarily in order to train faster):
  - CORnet-Z: 0.01, divided by 10 every 10 epochs;
  - CORnet-R: 0.1, divided by 10 every 10 epochs;
  - CORnet-S: 0.1, divided by 10 every 20 epochs.
- **Optimizer:** Stochastic Gradient Descent with momentum.9
- **Loss:** cross-entropy between image labels and model predictions (logits)
- **Code and weights:** github.com/dicarlolab/CORnet

## Benchmarking

### Brain-Score

(Schrimpf et al., 2018) is a composite benchmark that measures how well models can predict (a) the mean neural response of each neural recording site to each and every tested naturalistic image in non-human primate visual areas V4 and IT (data from Majaj et al., 2015) and (b) mean pooled human choices when reporting a target object to each and every tested naturalistic image (data from Rajalingham et al., 2018).

#### Neural predictability

A total of 2760 images containing a single object pasted randomly on a natural background were presented centrally to passively fixated monkeys for 100 ms and neural responses were obtained from 88 V4 sites and 168 IT sites. For our analyses, we used normalized time-averaged neural responses in the 70-170 ms window. A regression model was constructed for each neuron using 90% of image responses and tested on the remaining 10% in a 10-fold cross-validation strategy. The median over neurons of the Pearson’s *r* between the predicted and actual response constituted the final neural fit score for each visual area. In CORnets, we used designated model areas and the best time point to predict corresponding neural data. In comparison models, we used the most predictive layer.

#### Behavioral predictability

A total of 2400 images containing a single object pasted randomly on a natural background were presented to 1472 humans for 100 ms and they were asked to choose from two options which object they saw. 240 of those images with around 60 responses per object-distractor pair were used in further analyses, totalling in over three hundred thousand unique responses. For all models tested, we used the outputs of the layer just prior to transformation into 1000-value ImageNet-specific category vectors to construct a linear (logistic regression) decoder from model features. We used the regression’s probabilities for each class to compare model choices against actual human responses.

### Feedforward Simplicity

Given equally predictive models, we prefer a simpler one. However, there is no common metric to measure model simplicity. We considered several alternatives. One possibility was to use the total number of parameters (weights). However, it did not seem appropriate for our purposes. For instance, a single convolutional layer with many filter maps could have many parameters yet it seems much simpler than a multilayer branching structure, like the Inception block (Szegedy et al., 2017), that may have less parameters overall.

Moreover, our models are always tested on independent data sampled from different distributions than the train data. Thus, after training a model, all these parameters were fixed for the purposes of benchmarks, and the only free parameters are the ones introduced by the linear decoder that is trained on top of the frozen model’s parameters (see above for decoder details).

We also considered computing the total number of convolutional and fully-connected layers, but some models, like Inception, perform some convolutions in parallel, while others, like ResNeXt (Xie et al., 2017), group multiple convolutions into a single computation.

We therefore computed the number of convolutions and fully-connected layers along the longest path of information flow. For instance, the circuits in each of CORnet-S areas have the length of four since information is passed sequentially through four convolutional layers. Note that we counted recurrent (shared parameter) paths only once. If recurrent paths were counted the number of times information was passed through them, models with shared parameters would be no simpler than those using unique parameters at every time (i.e., feedforward models), which is counterintuitive to us.

We also wanted to emphasize that the difference between a path length of 5 and a 10 is much greater than between 105 and 110, so the path length was transformed by a natural logarithm. Finally, we wanted a measure of simplicity, not complexity, so the resulting value was inverted, resulting in the following formulation of Feedforward Simplicity,

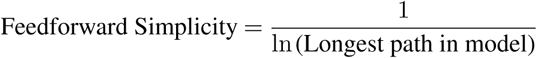

### Computability

All deep neural networks commonly used for predicting brain data fully specify how image pixels are transformed into high-level features. What remains unspecified, however, is at least the following:

- how these high-level features are transformed into behavioral choices and neural responses;
- which areas in the model correspond to which areas in the visual cortex.

These choices are left open to a researcher. Ideally, however, there would be no free parameters at all. CORnets attempt to make more commitments than other models by specifying which model areas correspond to which brain areas, making CORnet family more computable than alternatives. Importantly, this commitment also means that, for example, if model’s area V1 is worse than area V4 at predicting neural responses in visual area V1, then this model would be falsified.

## Results

### CORnet-Z is a lightweight AlexNet alternative

CORnetZ is very fast, achieving 48% ImageNet top-1 performance in less than 20 hours of training on a single Titan X. Despite worse performance than AlexNet (58%), it produces comparable V4, IT, and behavioral prediction scores with less computations (five convolutional and fully-connected layers versus eight; Fig. 3). (If desired, CORnet-Z’s ImageNet performance can be easily matched to AlexNet’s by adding an extra area after model’s IT area with 1024 output channels.) However, we observed that V4 data is predicted slightly better (by about .01) by model’s V2 area rather than V4. Since we committed to a particular model-to-brain mapping in CORnets, we use model’s area V4 for V4 neural prediction, but this mismatch already reveals a shortcoming in CORnet-Z.

**Fig. 3.**
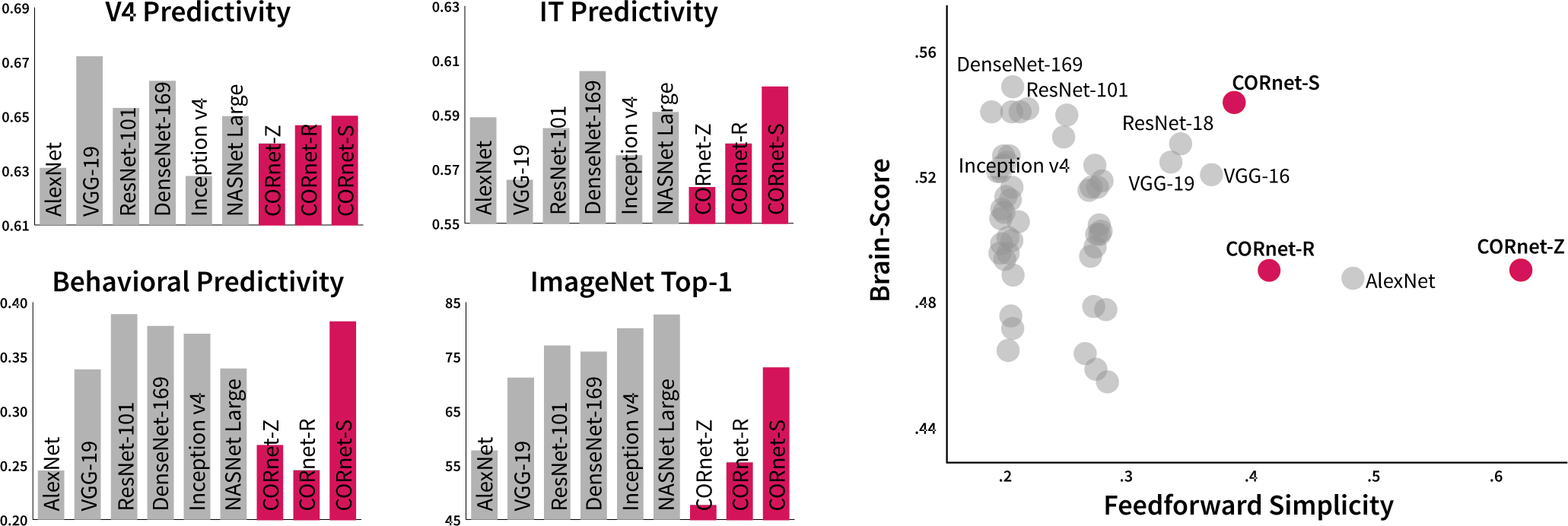
(*left*) Comparison of various scores on several popular models and three CORnet models. CORnet-Z and CORnet-R predict neural and behavioral responses close to AlexNet level, while CORnet-S is comparable to the state-of-the-art models. **(*right*) Feedforward Simplicity versus Brain-Score.** Most simple models perform poorly on Brain-Score, while best models for explaining brain data are complicated. CORnet-S offers the best of both worlds with the best Brain-Score and the highest degree of simplicity we could achieve to date. Note that dots were slightly jittered along the x-axis to improve visibility. Most of these jittered datapoints come from either MobileNet v1 (Feedforward Simplicity around .27) or v2 (Feedforward Simplicity around .2), so all models have the same simplicity (due to the same architecture) but varying Brain-Score (due to varying numbers of neurons).

### CORnet-R is a lightweight recurrent model

CORnet-R slightly improved neural fits over CORnet-Z and exhibited better behavioral predictability. It also improved in ImageNet top-1, presumably due to an extra convolution in each model’s area. Moreover, its recurrent nature allows for temporal predictions and analyses in dynamic tasks. However, similar to CORnet-Z, V4 neural responses were predicted slightly better by model’s V2 area rather than V4.

### CORnet-S is one of the best yet substantially simpler brain-predicting models

While all models in CORnet family are strong at neural prediction, CORnet-S also achieves a high behavioral prediction score, making it one of the best models tested on Brain-Score so far. Critically, CORnet-S is substantially simpler than other top-performing models on Brain-Score (Fig. 3, right) and commits to a particular mapping between model and brain areas.

However, note that CORnet family of models was developed using Brain-Score as a guiding benchmark and although it was never directly used in model search or optimization, testing CORnets on Brain-Score is not a completely independent test. Ideally, all models should be further validated on new experimental data sets – work that is in progress now.

## Discussion

### Is CORnet-S the correct model of Core Object Recognition?

No it is not. It is simply the model that best achieves our criteria at the moment. We believe that there are many other models that could achieve these criteria just as well or even better (e.g., Nayebi et al., 2018). As we find them, we will upgrade CORnet to the next version. Moreover, one should keep in mind that our current criteria are not necessarily the only or the “correct” criteria. Indeed, we expect our criteria to evolve as we acquire more data of the same kind or expand our datasets to include more images, tasks, and regions from which recordings or other data types are collected, or develop new methods for comparing models and brain data (see Discussion in Schrimpf et al., 2018).

### Are there really only four areas in the ventral visual stream?

While this is an experimental question that depends on one’s definition of “area”, we can say that four-area neural networks appear to be sufficient to reach state-of-the-art on Brain-Score. However, many researchers, including our own group, often divide IT into several sub-areas such as posterior IT (pIT), central IT (cIT), and anterior IT (aIT). While adding one more area in CORnet-Z, for instance, can result in a 10% increase in ImageNet top-1 performance, current brain-related metrics do not appear to be significantly affected by having more areas, thus we chose to use four areas for simplicity.

### Is it correct to assume that multiple convolutions can occur within a single cortical area?

Each neocortical area contains six cortical layers, with neurons in each layer potentially involved in different types of computations. Because neural types in each layer are replicated laterally across the cortical area, similar operations are thought to be applied across the area within each layer. Because the convolution operator is one way to model this, it is not unreasonable to assume that each cortical area includes the functional equivalent of multiple convolutions applied to the incoming sensory “image” transmitted from the previous area. Areas in CORnet-S can thus be thought to very crudely approximate the functions of the cortical microcircuitry considered at the level of the entire cortical area. For feedforward processing only, the minimum number of convolutions is two per area (one for layer IV and one for layer II/III) to create a new “image” transmitted to the next cortical area.

However, one must keep in mind that biological neurons themselves need not be characterized as performing a single weighted sum, followed by a non-linearity. Rather, such computations could be performed by multiple groups of dendrites, effectively reconceptualizing each biological neuron as a multilayer perceptron (Mel, 2016). Thus, it is currently difficulty to experimentally estimate how many convolutions (or convolution-like) operations are achieved by each cortical area, but the number enacted in each area of CORnet-S (namely, four) is not out of the range of possibilities.

## Limitations: What these CORnet models have not yet addressed

### Retina and LGN are lacking

So far all models we trained develop Gabor-like detectors in their first area. We call this first model area “V1”, which thus implies that our models do not have retina and LGN. One possibility is that given a different training strategy (e.g., unsupervised), retina-like and LGN-like functional properties would emerge in the first two layers of a naive model architecture. Alternatively, it is possible that retinal and LGN neurons have particular genetically-determined connectivity properties that cause their particular selectivities to develop and that we will be unable to obtain in model neural networks unless this particular circuitry is explicitly built into the model architecture. Whether explicitly adding retinal and LGN architectures in front of the model would alter model’s Brain-Score remains an interesting future research direction.

### Anatomy and circuitry are not precisely biomimetic

While our long term goal is a model of all the mechanisms of the ventral stream, we do not claim that CORnet models are precisely biomimetic. For example, we ignored that visual processing involves the retina and lateral geniculate nucleus (LGN) (see above). However, we believe that the CORnet models are closer approximations to the anatomy of the ventral visual stream than current state-of-the-art deep ANNs because we specifically limit the number of areas and we include some recurrence.

We note however, that many of our other model choices for these three CORnet models are driven by engineering requirements – most importantly, making sure the model’s non-architectural parameters can by set by gradient descent (i.e., the model can be trained) and that the model occupies few GPUs during train time. So, for instance, adding a skip connection was not informed by cortical circuit properties but rather proposed by He et al. (2016) as a means to alleviate the degradation problem in very deep architectures (where stacking more layers results in decreased performance). In CORnet-S, it appears to also have helped with a faster convergence on Brain-Score but we would be wary to claim biological significance. In that sense, it is possible that the circuit structures (a.k.a. areas) we selected are good for training with gradient descent on today’s hardware rather than being circuits that evolution converged upon.

On the other hand, we note that not just any architectural choices work. We have tested hundreds of architectures before finding these and thus it is possible that the proposed circuits could have a strong relation to biological implementations. Indeed, a high-level approach of searching for architectures given a suitable learning rule and task or guided by biological constraints may yield convergent circuits with biology (Bashivan et al., 2018).

Going forward building new CORnet models, our continuing goal will be to use models to ask which aspects of biological anatomy need to be better approximated or even mimicked by a CORnet model to achieve better functional matches to visual processing and COR behavior (i.e., better Brain-Scores). Our overarching logic is that ANNs that are most functionally similar to the brain will contain mechanisms that are most like those used by the brain.

### The model mechanisms that link from the neural processing to the object choice behavior are static

These linkage elements are referred to as the “decoder” portion of the model (see above). In the CORnet models used here, the decoder is a linear classifier, as it is one of the simplest classifiers that could be implemented in a biological neural network. However, the current decoder lacks dynamics and the ability to integrate evidence over time in interesting ways (Gold and Shadlen, 2007; O’Connell et al., 2018). For instance, for training recurrent CORnets we only use the last output of the IT layer. A more biologically-plausible decoder should at least be able to integrate information over time (see Kar et al., 2018b, for preliminary results).

### Learning mechanisms are non-biological

These COR-net models are only intended to be models of primate adult processing and behavior of 200 msec of image processing (the approximate duration of each natural primate fixation). We do not attempt to explain how the actual ventral visual stream reached this adult end state. At the moment, we assume an architecture (which might be delivered by evolution and pre-natal development), and we use supervised training and stochastic gradient descent (neither of which might be applicable to biological post-natal development and learning). However, this procedure is the only one known thus far to provide a reasonably accurate quantitative model of adult mid- and high-level neural responses and behavioral responses and therefore may be a suitable supervised proxy to arrive at models that are similar to those obtained in the future by more biologically-plausible learning mechanisms (e.g., self-supervised tasks and local learning rules). This is clearly an important and wide open area of future work.

## ACKNOWLEDGEMENTS

We thank Pouya Bashivan and Kohitij Kar for useful discussions and insights, and Maryann Rui and Harry Bleyan for the initial prototyping of the CORnet family. We also thank Ricardo Henriques for this bioRxiv LATEX template.

This project has received funding from the European Union’s Horizon 2020 research and innovation programme under grant agreement No 705498 (J.K.), US National Eye Institute (R01-EY014970, J.J.D.), Office of Naval Research (MURI-114407, J.J.D), the Simons Foundation (SCGB [325500, 542965], J.J.D; 543061, D.L.K.Y), the James S. McDonnell foundation (220020469, D.L.K.Y.) and the US National Science Foundation (iis-ri1703161, D.L.K.Y.). This work was also supported in part by the Semiconductor Research Corporation (SRC) and DARPA.

